# “Cell surface associated LapA of *Pseudomonas fluorescens* is anchored inside its Type-1 Secretion TolC-like Pore”

**DOI:** 10.1101/198937

**Authors:** T. Jarrod Smith, Holger Sondermann, George A. O’Toole

## Abstract

The type-1 secretion system (T1SS) of gram-negative bacteria enables a one-step translocation strategy known to move functionally diverse proteins from the cytoplasm into the extracellular environment without a periplasmic intermediate. LapA of *Pseudomonas fluorescens* Pf0-1 is a giant type-1 secreted (T1S) adhesin that facilitates biofilm formation only when displayed at the cell surface. A LapA-targeting periplasmic protease, LapG, connects intracellular cyclic diguanylate (c-di-GMP) levels with cell surface-associated LapA by cleaving and absolving LapA from the cell surface under conditions unsuitable for biofilm formation. Here, we demonstrate that LapA contains a novel N-terminal element, called the retention module (RM), which prohibits classical one-step T1S of LapA. We provide evidence that the RM of LapA tethers LapA at the cell surface through its outer membrane TolC-like pore, LapE, where LapA is accessible to the periplasmic protease LapG. We also demonstrate that this unusual retention strategy is likely conserved among LapA-like proteins and represents a new subclass of T1SS ABC transporters exclusively involved in transporting LapA-like adhesins.

**Significance Statement:** Bacteria have evolved multiple secretion strategies to interact with their environment. For many bacteria, the secretion of cell surface associated adhesins is often key for initiating contact with a preferred substrate to facilitate biofilm formation. Our work demonstrates that *P. fluorescens* uses a previously unrecognized secretion strategy to retain the giant adhesin LapA at its cell surface. Further, we identify likely LapA-like adhesins in various pathogenic and commensal Proteobacteria and provide phylogenetic evidence these adhesins are secreted by a new subclass of T1SS ABC transporters.

## Text

The biofilm lifestyle is profoundly consequential to human health and industry, for better and worse (1, 2). Although bacteria initiate surface contact and biofilm formation through a variety of strategies, many microbes need cell surface-associated protein adhesins to bind a surface. The family of giant (>200 kDa) T1S repeats-in-toxin (RTX)-containing adhesins is known to be critical for biofilm formation or surface binding by a variety of organisms, including *Pseudomonas*, *Bordetella*, *Legionella*, *Vibrio, Shewanella* and *Marinomonas* (3–8).

Mechanistic studies from our lab on the giant RTX adhesin, LapA, of *Pseudomonas fluorescens* Pf0-1 identified a novel regulatory node, LapDG, that links cell surface associated LapA levels to intracellular c-di-GMP levels. Here, the activity of the LapA-targeting periplasmic protease, LapG, is inhibited when its effector, LapD, a transmembrane protein, is bound to cellular c-di-GMP. Conversely, when c-di-GMP levels decrease in the cell, LapD releases LapG. Free LapG in turn cleaves LapA from the cell surface, releasing the adhesin into the supernatant where it is inoperative for biofilm formation (9, 10). Homologs of the *lapDG* genes are found through the Proteobacteria, suggesting that this poorly understood surface display strategy may be quite common (11, 12).

Like other T1S proteins, such as HlyB of *E. coli* and CyaA of *B. bronchiseptica*, LapA’s C-terminal secretion signal and cognate T1S machinery (LapEBC) are required for LapA secretion and thus biofilm formation (6, 13). T1SS is considered a one-step secretion strategy that lacks any periplasmic intermediate (14), and it is currently unclear how LapA localizes to the cell surface with an N-terminal dialanine cleavage motif that is accessible to the periplasmic protease, LapG. Here, we demonstrate that LapA is not secreted in a canonical one-step T1SS fashion, but rather tethers to the cell surface through its T1SS apparatus. A cleavable retention module at the N-terminus of LapA prohibits complete secretion until cleaved by the LapG protease. We also provide evidence that this previously unappreciated retention strategy is broadly conserved in Proteobacteria and represents a third, distinct subclass of T1S systems.

## Results

### Bioinformatic Identification of LapG Substrates

Despite the importance of LapA and related proteins as key biofilm adhesins (15), it is difficult to identify LapG substrates due to their relatively low sequence similarly. Additionally, ORF analysis programs often overlook or misannotate these large and complex adhesins (5, 16, 17). To overcome the first limitation, we took advantage of the observation that the two proteins that control LapA localization, the c-di-GMP-receptor LapD and the LapD-regulated protease LapG, show high sequence similarity and functional conservation between microbes (3, 12, 18) and can be identified by their respective domain architectures (LapG, pfam06035; LapD, pfam16448). We utilized the NCBI conserved domain database (CDD) and genome database to identify bacterial species encoding *lapDG* homologs; ∼1300 such *lapGD*-encoding species spanning 120 genera were identified. Each annotated genome was investigated for proteins containing hallmarks of LapA: an N-terminal LapG cleavage site and C-terminal RTX motifs. To accomplish this task, we developed an algorithm to recognize large proteins (>1000 aa) with RTX-motifs (Dx[L/I]x(4)GxDx[L/I]xGGx(3)D) and a canonical N-terminal dialanine LapG cleavage motif ([T/A/P]AA[G/V]). Although our approach is constrained to properly annotated LapA-like ORFs, we still identified over 500 putative LapG substrates in ∼50 genera throughout Proteobacteria, including *Legionella* and *Vibrio* species (Table S1). Importantly, characterized LapG substrates LapA and BrtA (3, 19) were identified. Interestingly, some species likely encode multiple LapG substrates, including *P. fluorescens* Pf0-1(LapA and Pfl01_1463) and *V. cholerae* O395 (FrhA and VC0395_0388).

### LapG Substrates Predicted *in silico* Are Processed *in vitro*

LapG homologs cleave a variety of LapA-like N-termini from unrelated species in *vitro* and *in vivo* (3, 18). To help validate the utility of our algorithm and confirm predicted LapG substrates, we cloned and expressed the N-terminal elements (∼250-350 aa) of the putative LapG-proteolyzed adhesins Pfl01_1463 from *P. fluorescens* and VC0395_0388 from *Vibrio cholerae* (Fig 1A). Cell lysates of *E. coli* expressing C-terminally 6HIS-tagged, N-terminal fragments of Pfl01_1463 (M1-240S, 117TAAG120) or VC0395_0388 (M1-363G, 127AAAG130) were mixed with a lysate made from *E. coli* expressing *P. fluorescens* Pf0-1 LapG from a plasmid or the empty vector control. The LapG-dependent cleavage product was tracked via Western blot, with the N-terminus of LapA (M1-235V, 107TAAG110) and an uncleavable variant (LapA^TRRG^, AA108-109RR) serving as positive and negative controls, respectively (9). *P. fluorescens* Pf0-1 LapG cleaves both predicted substrates (Fig 1 B & C). Unlike FrhA (4), a role for VC0395_0388 in infection and/or biofilm formation is currently unknown; however, these data implicate VC0395_0388 as a cell-surface associated, c-di-GMP-regulated biofilm-promoting adhesin.

**Figure 1.**
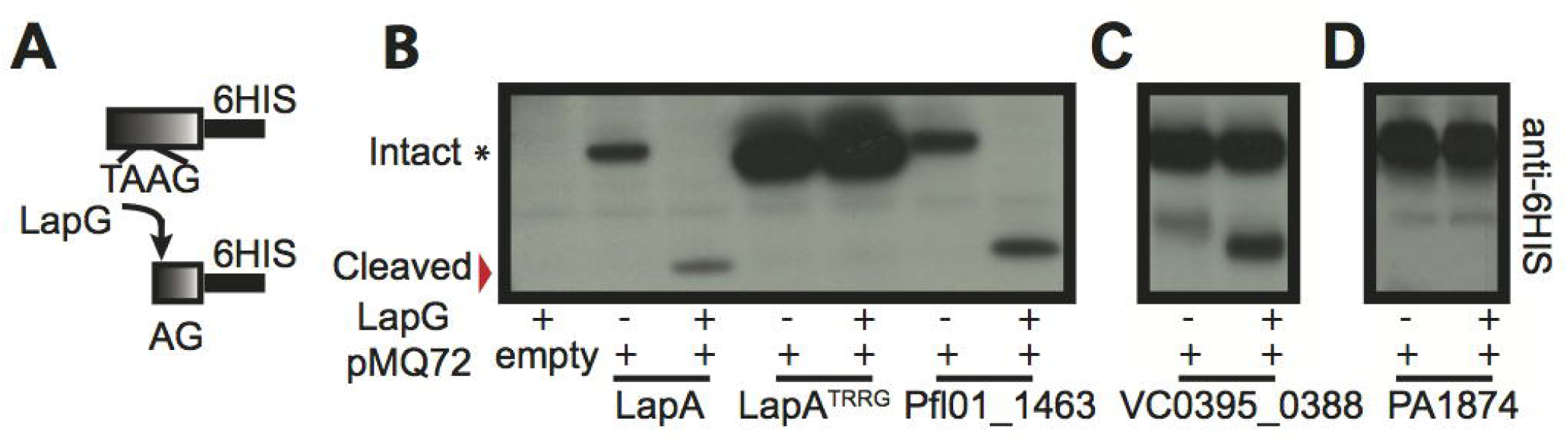
*P. fluorescens* Pf0-1 LapG *in vitro* cleavage analysis. (A) Overview of *in vitro* LapG cleavage assay. (B-D). Intact (*) and cleaved (red arrow) N-termini were visualized by Western blot. Equal protein concentrations of the substrate and extracts from LapG-expressing strains were used.

Conversely, *P. fluorescens* Pf0-1 LapG is unable to cleave the N-terminus of PA1874(M1-251T, 137AAAIG141) (Fig 1D), a large 238 kDa putative outer membrane adhesin encoded by *Pseudomonas aeruginosa* PAO1. Although PA1874 resembles LapA, it was not detected by our algorithm because it contains a degenerate LapG cleavage motif (137AAAIG141). Instead, we manually identified PA1874 by its proximity to *lapEBC* homologs (20). Together, these data support the predictive power of our algorithm for detecting LapA-like LapG substrates *in silico*, and demonstrate the breadth of this c-di-GMP-regulated biofilm strategy in Proteobacteria.

### *P. fluorescens* LapA Homologs Contain a Conserved N-terminal Domain

Previous work from our lab implicated the N-terminus of LapA in retaining the giant adhesin at the cell surface (13, 19). To gain insight into LapA’s retention mechanism, we aligned putative LapA-like proteins from closely related *P. fluorescens* strains detected *in silico* and color-filled residues according to the default Clustal X color parameters in Jalview to highlight highly similar regions (Fig S1A, Table S2). Our analysis revealed these adhesins contain two highly similar regions: a C-terminal region that corresponds to LapA’s T1S signal required for secretion (Fig S1A, blue box) and a N-terminal region that extends ∼20 residues beyond the LapG dialanine cleavage motif (Fig S1A, red box and Fig 2A). Given the C-to N-terminal secretion directionality of T1S substrates (21), we speculated that the N-terminus may be involved in retaining these giant adhesins at the cell surface. Consistent with this idea, analysis of BrtA and RtxA homologues from *B. bronchiseptica* and *L. pneumophila* strains, respectively, revealed similar patterns of high identity N- and C- termini (Fig S1B,C).

**Figure 2.**
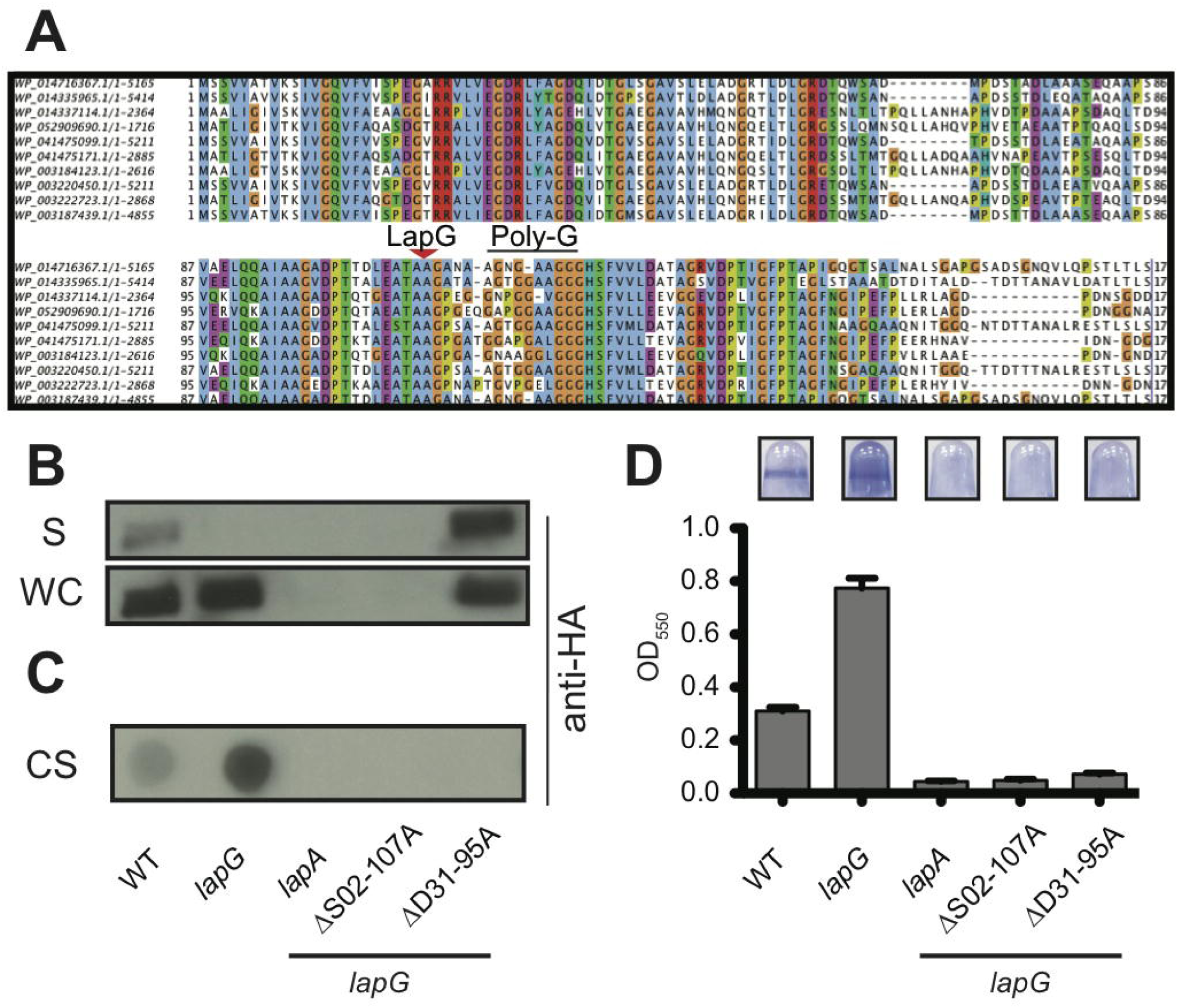
LapA’s N-terminus Serves as a Retention Module. (A) The first ∼175 aa of the *P. fluorescens* LapA-like N-termini (Fig. S1A, red box) with LapG cleavage motif (red arrow) and putative polyglycine linker (Poly-G, black bar). (B) Western blot analysis of supernatant and whole cell fractions for LapA. (C) Dot blot analysis of surface-associated LapA. (D) Biofilm analysis of LapA N-terminal mutants (n=8, +/-SEM).

### The N-terminus of LapA is Required for Surface Retention

Closer examination of the N-terminus of *P. fluorescens* LapA homologs indicates their sequence similarity breaks down shortly after a poly-glycine region (Figure 2A, Poly-G). This poly-glycine region is present in many of the LapA-like proteins predicted by our algorithm (Fig S2A). Because poly-glycine regions often serve as unstructured domain linkers, we hypothesized the N-terminal region of LapA encompassing up to this linker may function as a retention module (M1-125S).

To test this idea, we made targeted deletions in this putative N-terminal domain. Analysis of the primary sequences and predicted secondary structures suggests the N-termini of the LapA-like proteins identified by our algorithm share little sequence identity, but may adopt similar secondary structures (Fig S2A-B). Given that glycine residues are known to punctuate secondary structures, we used a “gly-to-gly” targeted truncation strategy to disrupt secondary structures within the N-terminus of LapA, using the alignment from Fig S2A as guide.

LapA is typically undetectable in the supernatant and enriched at the cell surface of the *lapG* mutant (Fig 2B and 2C, WT vs *lapG*). Therefore, RM mutants were engineered into the hyper-biofilm forming *lapG* gene deletion strain using unmarked allelic replacement, allowing us to decouple retention defects from LapG-mediated proteolysis. Biofilm formation and LapA localization were assayed by comparing each retention mutant to the parental *lapG* mutant, and Western blotting for LapA in the whole cell (WC), supernatant (S) and cell surface (CS) fractions.

Consistent with our hypothesis, a LapA truncation mutant lacking residues D31-95A (ΔD31-95A) is unable to form a biofilm (Fig 2D), and this LapA variant is found in the supernatant (Fig 2B) but not retained at the cell surface (Fig 2C). While the ΔV23-95A and ΔD31-95A phenotypes were indistinguishable, most mutants engineered for this study showed only a slightly reduced biofilm phenotype (Fig S2B-C). For those mutants with a biofilm defect (i.e., ΔS02-107A), Western analysis showed that the mutant protein was unstable. Together, these data indicate the N-terminus of LapA functions as a retention module, contributing to the localization of LapA to the cell surface, and also suggests that this region plays a critical, but unclear, role in LapA stability.

### Cell Surface Associated LapA Engages the LapEBC T1SS Machinery Through its N-Terminal Domain

Next, we investigated how the RM tethers LapA to the cell surface. LapA is found in the outer membrane fraction (24); however, LapA’s RM lacks elements previously shown to be involved in forming outer membrane pores and translocation structures in target cell membranes (15, 22, 23). Thus, we considered the possibility that LapA is anchored to the cell surface by remaining in the T1SS apparatus as a secretion intermediate, using its N-terminal RM to prevent complete secretion of the adhesion. Although unheard of for T1S proteins, we previously demonstrated the LapG substrate from *Pseudomonas aeruginosa*, CdrA, a two partner secreted protein, uses a “cysteine-hook” formed by an intramolecular disulfide bond to anchor itself to the cell surface through its outer membrane pore, CdrB (12, 25). Additionally, recent structural analysis and modeling performed on MpIBP, a large LapA-like adhesion of *M. primoryensis*, suggests the RM of MpIBP, which shares secondary structure features with LapA, may form a plug that prohibits complete secretion of the adhesin through its outer membrane T1S-associated TolC-like protein (26). However, to date, there has been no experimental data supporting the retention model we explore here.

To determine if LapA is secreted via the classical one-step T1SS model or retained within its translocation machinery, we conducted a secretion competition experiment to compare secretion of the C-terminal secretion domain of LapA tagged with a HA-epitope tag for Western blotting (HA-C235; Fig 3A, far right). This tool allows us to discern if cell surface-associated LapA impacts the availability of LapEBC T1SS to secrete the HA-C235 peptide. We examined secretion of the HA-C235 protein in strains where LapA is locked at the cell surface (a Δ*lapG* mutant), or alternatively, in strains where LapA is continuously secreted into the supernatant (Δ*lapD* or *lapG lapAΔD31-95A* mutants). In the Δ*lapD* mutant, constitutive LapG activity removes LapA from the cell surface, while the *lapG lapAΔD31-95A* strain expresses a variant of LapA that lacks the complete retention signal (see Fig 2).

**Figure 3.**
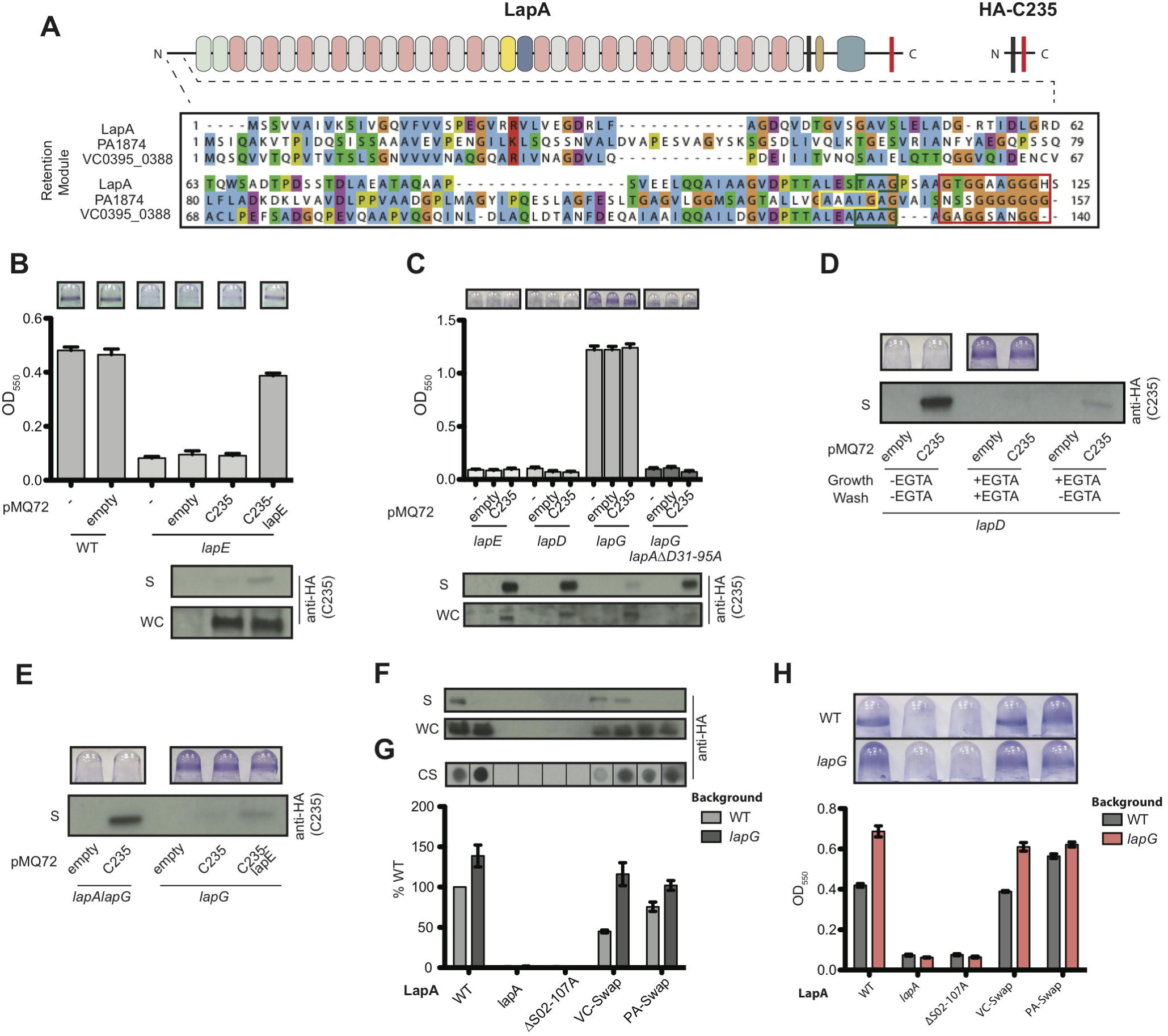
LapA Cell Surface Localization Impacts LapEBC Activity. (A) Top: Scaled representation of LapA and HA-C235. Black bar indicates 3XHA tag in LapA and HA-C235. Bottom: MUSCLE alignment of putative RMs from *P. fluorescens*, *P. aeruginosa*, and *V. cholerae* adhesins with residues colored according to the default Clustal X coloring scheme in Jalview. The putative polyglycine linker is boxed in red while the canonical and degenerate LapG cleavage sites are boxed in green and yellow, respectively. (B-C) Biofilm formation (top) and Western blot analysis of HA-C235 in the supernatant (S) and whole cell (WC) fractions (bottom). In these experiments, LapE and HA-C235 are co-expressed as a transcriptional fusion from a plasmid. (D) Biofilm analysis (top) and HA-C235 secretion (bottom) in Δ*lapD* mutants with and without EGTA treatment. (E) Biofilm analysis (top) and HA-C235 secretion (bottom) in Δ*lapG* mutants. (F) Western blot analysis of LapA RM chimeras VC0395_0388 (VC-Swap; also panel A, bottom.) and PA1874 (PA-Swap; also panel A, bottom) in the supernatant (S) and whole cell (WC) fractions in the indicated genetic backgrounds. (G) Western blot analysis of cell surface associated LapA RM chimeras (top) and quantification (bottom). Strains in panel G correspond to strains in panel F. (H) Biofilm formation of LapA RM chimeras from strains in panel G. For biofilm and dot blot quantification (n=8, +/-SEM).

Western blot analysis of the supernatant fraction indicates like LapA, HA-C235 peptide secretion is LapEBC–dependent. Deletion of *lapE* results in loss of biofilm formation, as reported previously (6) and eliminates HA-C235 secretion (Fig 3B). HA-C235 secretion is restored when HA-C235 and *lapE* are introduced into the Δ*lapE* mutant and expressed as a transcriptional fusion from a plasmid (Fig 3B, far right).

In the Δ*lapA*Δ*lapG* mutant, HA-C235 is constitutively secreted because it lacks the LapA RM (Fig 3C). HA-C235 supernatant levels in the Δ*lapA*Δ*lapG* and Δ*lapD* mutants are identical, indicating that lack of LapA or constitutive LapA secretion, respectively, does not impinge on HA-C235 secretion (Fig 3C, left). However, when LapA is locked at the cell surface in the Δ*lapG* mutant strain, HA-C235 secretion is nearly abolished (Fig 3C, center). HA-C235 secretion is restored in a Δ*lapG* mutant when LapA does not associate with the cell surface (compare Δ*lapG* mutant to Δ*lapG lapAΔD31-95A*; Fig 3C). Together, these data indicate retention of LapA, but not its secretion, limits LapEBC availability for secreting other peptides, and is consistent with the model that the RM of LapA is retained in the LapEBC secretion apparatus.

### Chemical Inhibition of LapG Locks LapA at Cell Surface, Occluding the LapEBC T1SS Machinery

LapG is a calcium-dependent cysteine protease that can be chemically inhibited *in vivo* and *in vitro* with micromolar levels of EGTA (19). Addition of EGTA to the Δ*lapD* mutant, which usually exhibits constitutive LapG activity, restores LapA cell-surface localization by inhibiting LapG (19). To determine if chemical inhibition of LapG in the Δ*lapD* background can block HA-C235 secretion by enhancing surface associated LapA, we compared biofilm formation and HA-C235 secretion in the Δ*lapD* mutant grown in the absence or presence of 500 µM EGTA (-EGTA or +EGTA). Each strain was grown for 5 hr, then washed and resuspended for 20 min in the indicated medium (Fig 3D, growth/wash). The supernatants were then collected and HA-C235 protein was probed via Western blot analysis.

EGTA treatment restored biofilm formation and inhibited HA-C235 secretion in the Δ*lapD* mutant (Fig 3D). Additionally, washing EGTA grown cells in medium lacking EGTA restored C235 secretion (Fig 3D, right), indicating that removal of EGTA reactivated LapG activity, allowing this protease to cleave LapA from the cell surface and thus allow secretion of HA-C235. These data support our model that LapA localizes to the cell surface via anchoring in its T1S machinery.

### LapE Overexpression Rescues HA-C235 Secretion in a *lapG* Mutant Background

The LapEBC proteins likely form a tripartite complex (LapE-LapBC) to secrete LapA. Given that secretion of HA-C235 is blocked when LapA is locked at the cell surface (i.e., a *lapG* mutant or addition of EGTA), we reasoned that a T1SS component(s) participating in cell surface localization of LapA is likely limiting. Therefore, overexpression of the limiting secretion component should allow additional secretion, and thus should rescue HA-C235 secretion in a Δ*lapG* mutant. Previous work from our lab comparing *lapA, lapE, lapB*, and *lapC* gene expression in the presence and absence of the biofilm promoting nutrient phosphate indicates only *lapE* is down regulated under phosphate-limiting conditions that inhibit biofilm formation (27). Thus, we suspected that *LapE* might be the limiting component of the secretion machinery.

To determine if *lapE* overexpression can rescue HA-C235 secretion in a *lapG* mutant, we assayed for HA-C235 in the supernatant of the Δ*lapG* mutant carrying a pC235-*lapE* transcriptional fusion construct. This construct allows simultaneous expression of HA-C235 and LapE protein from the same promoter. Western analysis of the supernatant fraction indicates co-expression of HA-C235 and LapE, but not HA-C235 alone, can rescue HA-C235 secretion in a *lapG* mutant background (Fig 3E, right). HA-C235 secretion in the Δ*lapA*Δ*lapG* mutant was used as a baseline secretion control. Together, these data are consistent with the model that LapA is retained on the cell surface by LapE, LapE levels are limiting, and retained LapA occupies the secretion pathway thus blocking secretion of any other substrate.

### Evidence of a Conserved Retention Strategy

PHYRE analysis (28) predicts the N-termini of putative LapG substrates may adopt similar secondary structures, suggesting these adhesins could tether through the outer membrane like LapA. To test this idea, we replaced the RM of LapA (M1-125S) with low-identity putative RMs from *V. cholerae* and *P. aeruginosa* proteins described earlier (Fig 3F,G, also Fig 1C,D). Figure 3A shows the LapG cleavage site in LapA and VC0395_0338 (boxed in green), the degenerate LapG cleavage site in PA1874 (boxed in yellow), and the putative polyglycine linker common to all three N-termini (boxed in red). Based on our *in vitro* LapG cleavage experiments, the putative RMs may represent separate classes of LapA-like adhesins. VC0395_0338 has a conserved LapG cleavage motif (AAAG) and is processed by LapG *in vitro* (Fig 1C). In contrast, while the N-terminus of the *P. aeruginosa* protein PA1874 has features similar to LapA, it appears to contain a degenerate LapG processing motif (AAAIG) and is not processed in vitro (Fig. 1D). Thus, we predicted PA1874 anchors to the cell surface but is not subject to LapG-mediated proteolysis and release. Using allelic exchange, we replaced the DNA encoding the N-terminus of LapA with DNA corresponding to the amino acids detailed in the alignment in Fig 3A in both the wild type *P. fluorescens* and the Δ*lapG* mutant background. These chimeras are called VC-Swap and PA-Swap. Biofilm formation and chimera localization were compared for the wild type and the Δ*lapG* mutant strains expressing the chimeras to determine if these proteins were retained on the surface, could support biofilm formation in *P. fluorescens* and/or were subject to LapG proteolysis to release the chimeric adhesins from the cell surface.

Western blot analysis of the whole-cell fraction indicates both VC-Swap and PA-Swap chimeras are stable in the wild type (light grey) and *lapG* (dark grey) mutant backgrounds (Fig 3F, WC). The biofilm phenotype and chimera localization of the VC-Swap and PA-Swap retention chimeras are consistent with their differential susceptibility to LapG cleavage *in vitro* (Fig 1C and D). Western blot analysis of the VC-Swap chimera in the supernatant (Fig 3F, S) and cell surface fractions (Fig 3G, CS) indicates deletion of the *lapG* gene decreases levels of the chimera in the supernatant fraction (Fig 3F), and more importantly, enhances chimera levels at the cell surface (Fig 3F, G; VC-Swap, light grey vs. dark grey) and increased biofilm formation (Fig 3H, VC-Swap; light grey vs. red bar). This trend mimics wild type LapA in the parental wild type and *lapG* backgrounds (Fig 3H WT; light grey vs. red bar). These data indicate the RM from VC0395_0338 (Fig 3A, bottom) can complement LapA localization and LapG-dependent release from the cell surface, suggesting LapA and VC0395_0338 localize to the cell surface by a similar mechanism. Conversely, PA-Swap chimera localization and corresponding biofilm formation were not impacted by LapG activity. Consistent with our *in vitro* LapG cleavage analysis, these data indicate that the PA1874 RM can complement LapA’s RM for cell surface retention and biofilm formation, but not for cleavage by LapG due the presence of a degenerate cleavage motif (Fig 3F-H).

### ABC Transporters of LapA-like Adhesins Form a Distinct T1SS Subgroup

ABC transporters are found throughout all three domains of life, but can often be functionally grouped based on common residues that are critical for secreting their substrate(s) (29– 31). Studies of the T1SS ATPase HlyB of *E*. *coli*, which transports the RTX toxin HlyA, demonstrated HlyB contains a N-terminal domain that is critical for binding HlyA’s C-terminal RTX motifs (31). The N-terminal domain of HlyB resembles the C39 peptidase domain (C39) typically found at the N-terminus of the ATPase component of bacteriocin ABC transporters, but HlyB lacks the catalytic cysteine residue required to cleave and activate immature bacteriocins during secretion (32). Instead, the C39-like domain (CLD) of HlyB contains a tryptophan involved in binding HlyA that is conserved among many ABC transporters involved in secreting RTX toxins, including the ABC transporter secreting CyaB of *B. bronchiseptica*. Thus, the C39 and CLD of bacteriocin and RTX toxin transporters can be distinguished by their amino acid sequence (31).

Interestingly, the T1SS ATPase for LapA, called LapB, contains a N-terminal domain that lacks both the conserved cysteine and tryptophan residues of C39 and CLD involved in bacteriocin-processing and RTX-binding, respectively. Given LapA’s unusual retention strategy and the observation that LapEBC is often encoded nearby LapDG homologs (11, 33), we were curious if LapB-like transporters may represent a distinct subgroup of ABC transporters involved in secreting adhesins with the newly defined N-terminal RM. To test this idea, we assessed the phylogenetic relationship between C39 and CLD sequences from the ATPases of bacteriocin and RTX toxin transporters, and the N-terminal domain of the ATPase from transporters involved in adhesin retention. The phylogenetic relationships of N-terminal domains from the ATPase component several characterized RTX toxin and bacteriocin transporters were identified using Interpro (http://www.ebi.ac.uk/interpro/), and compared with N-terminal domain of the putative LapB-like ATPase component.

LapB-like ABC transporters were defined as those encoded near LapDG homologs. We also included LssB of *L. pneumophila* and PA1876 of *P. aeruginosa* in the analysis. Recent studies indicate LssB is an ABC transporter involved with secreting an RTX adhesin, RtxA, which was predicted to be a LapG substrate by our algorithm (Fig S1B) (34). PA1876 is likely involved with transporting PA1874, the LapA-like protein with a degenerate cleavage motif described here (Fig 1D and Fig 3F-H) (20).

Our phylogenic analysis suggests the ATPase component of the ABC transporters of LapA-like adhesins form a distinct group that lack the functional residues critical for RTX motif-binding and bacteriocin-processing (Fig 4A, see WebLogo). These differences at the amino acid level likely reflect functional differences, rather than phylogenetic diversity of the organisms analyzed, because characterized RTX toxin and LapA-like adhesin ATPases encoded within the same genome cluster with their predicted substrate type. For example, the RTX toxin transporter CyaB (Fig 4A; orange circle) and LapA-like adhesin transporter BB1189 (red circle) encoded by *B. bronchiseptica* map to different clusters in this analysis. These data support the idea that adhesins with an N-terminal RM are secreted by a distinct group of T1SS ABC transporters.

**Figure 4.**
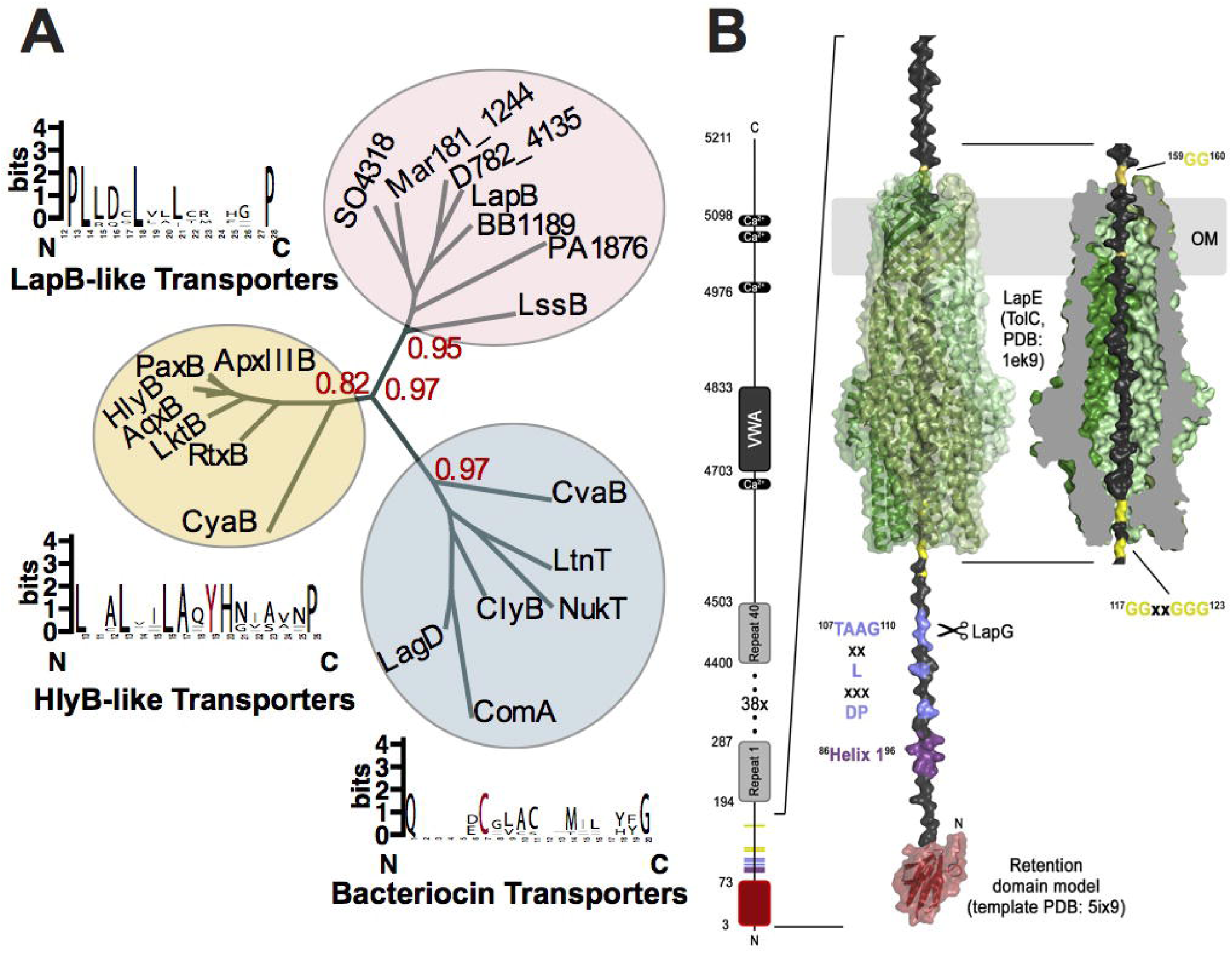
Phylogenetic Analysis of ABC Transporter Subfamilies. (A) Phylogenetic analysis of C39 peptidase domain, C39-like peptidase domain (CLD), and N-terminal ∼125 aa of LapB-like ATPases (Table S4) using the online phylogenetic analysis program phylogeny.fr under default settings (http://www.phylogeny.fr). WebLogos were generated by MUSCLE aligning members from each branch and truncated to highlight functional residues when applicable (red). Bootstrap values are indicated in red. (B) Left: Domain architecture of LapA. Right: Surface retention model for LapA. The structural model of the LapA retention module was generated using the Phyre2 protein fold recognition server and PDB 5ix9 as the template (red domain). The sequence following the retention domain was modeled as an extended peptide to reflect secondary structure predictions indicating this region (residues 97-173) to be flexible. The unstructured sequence was threaded through LapE, represented by the crystal structure of TolC (PDB: 1ek9). The LapG-targeting dialanine motif (TAAG) and regions critical for LapG binding (Helix 1 and DPxxxLxx) are shown, and proposed to be exposed in the periplasm. Two glycine-rich motifs in LapA, which flank the sequences within the TolC pore, are indicated in yellow.

### Discussion

Here, we describe a novel N-terminal element in LapA - called the retention module - that is responsible for localizing this giant adhesin to the cell surface. We propose structural features within LapA’s RM play a role in tethering LapA to the cell surface and show that disrupting these predicted secondary structures can lead to uncontrolled release of the adhesin from the cell surface, as was observed for the LapAΔD31-95A mutant. Consistent with these observations, low sequence identity RMs from putative LapA-like proteins share similar predicted secondary structures and can complement LapA function and localization, suggesting LapA’s unusual retention strategy is broadly conserved among this group of adhesins. The bioinformatic analysis presented here and previously (33, 35) suggest LapA-like proteins are secreted by an unappreciated subgroup of T1SS ABC transporters that engage TolC-like pores dedicated to the secretion and retention of these large adhesins.

How does the RM contribute to LapA’s cell surface localization? We propose the RM stalls the final steps of LapA translocation, leaving LapA threaded through the TolC-like outer membrane pore, LapE, with its RM localized in the periplasm, accessible to LapG, and C-terminal adhesive repeats displayed at the cell surface (Fig 4B). Although this model opposes the classical one-step paradigm detailed in over ∼25 years, the model is consistent with our artificial HA-C235 substrate competition experiments, EGTA-mediated inhibition of LapG activity, and expression studies with the TolC-like LapE outer membrane protein presented here. Given the abundance of LapA-like adhesins in pathogens and environmental microbes, and the conserved role of c-di-GMP in regulating their cell surface localization, this means of secreting and anchoring an adhesion appears to be a general strategy used for cell-surface and perhaps cell-cell adherence.

Based on the recently reported *Marinomonas* MpIBP structural modeling (26), and the data presented here and reported previously (13), we built a model of the LapA retention module (Fig 4B). Our genetic analysis of the N-terminal RM is in good agreement with the proposed N-terminal domain of MpIBP being localized to the periplasm. Interestingly, the proposed model for the MpIBP adhesin of *Marinomonas* (26) indicates the LapG proteolysis site may be obscured by the TolC pore, and our mutational analysis suggests regions not included in the MpIBP NMR structure are also important for retention and LapG processing (13). In contrast, we propose that the LapG processing site is accessible in the periplasm, a conclusion consistent with our modeling (Fig. 4B) and our previous biochemical and genetic studies (9, 12, 24).

## Materials and Methods

### Strains and media

*P. fluorescens* and *E. coli* strains listed in Table S3 were grown on lysogeny broth at 30°C and 37°C, respectively. Gentamycin was used when appropriate (10 µg/mL for *E. coli*, 30µg/mL for *P. fluorescens)*. For biofilm and LapA localization analysis, *P. fluorescens* strains subcultured in K10T-1 for 6 hr statically and with rotation, respectively (27).

### Static Biofilm Assay

The static biofilm assay was performed and quantified as described previously (13).

### LapA Localization

Whole cell, supernatant, and cell surface localization analysis was preformed as previously using a HA-tagged variant of LapA (13).

### HA-C235 Localization

Each strain was subcultured for 5.5 hr, washed, and then resuspended for 20 min in the indicated medium under non-inducing conditions. For chemical inhibition assays, K10T-1 medium was supplemented with 500µM EGTA as noted. Whole cell and supernatant fractions were prepared as for LapA localization. Western blot analysis against the HA epitope was used to detect HA-C235.

### *In Vitro* LapG Cleavage Analysis

*in vitro* cleavage analysis was performed as described previously (13).

### *In Silico* Prediction of LapG Substrates

LapG and LapD homologs were defined as ORFs encoding proteins with the pfam06035 and pfam16448 domains, respectively. The NCBI CDD was utilized to generate a list of LapG- and LapD-encoding bacteria, and the programming language R was used to determine the intersecting LapD/LapG-encoding bacteria. The protein annotation of these genomes were downloaded from the NCBI genome database and each annotated locus was interrogated for the presence of a LapG cleavage site within amino acids 80-150 ([T/A/P]AA[G/V]) and at least one RTX motif (Dx[L/I]x(4)GxDx[L/I]xGGx(3)D).

## Acknowledgment

We thank C. Boyd for building one of the control strains used in this study, T. Silhavy for his insight interpreting some of the data presented here.

